# Using Neural Networks to Improve Single Cell RNA-Seq Data Analysis

**DOI:** 10.1101/129759

**Authors:** Chieh Lin, Siddhartha Jain, Hannah Kim, Ziv Bar-Joseph

## Abstract

While only recently developed, the ability to profile expression data in single cells (scRNA-Seq) has already led to several important studies and findings. However, this technology has also raised several new computational challenges including questions related to handling the noisy and sometimes incomplete data, how to identify unique group of cells in such experiments and how to determine the state or function of specific cells based on their expression profile. To address these issues we develop and test a method based on neural networks (NN) for the analysis and retrieval of single cell RNA-Seq data. We tested various NN architectures, some biologically motivated, and used these to obtain a reduced dimension representation of the single cell expression data. We show that the NN method improves upon prior methods in both, the ability to correctly group cells in experiments not used in the training and the ability to correctly infer cell type or state by querying a database of tens of thousands of single cell profiles. Such database queries (which can be performed using our web server) will enable researchers to better characterize cells when analyzing heterogeneous scRNA-Seq samples.

Supporting website: <u>http://sb.cs.cmu.edu/scnn/</u>

Password for accessing the retrieval task webserver: scRNA-Seq

## 1. Introduction

Single cell RNA-seq (scRNA-seq) which profiles the transcriptome of individual cells (as opposed to ensemble of cells) has already led to several new and interesting findings. These include the level of heterogeneity within a population of cells [5], the identification of new markers for specific types of cells [19], and the temporal stages involved in the progression of various developmental processes [41].

While promising, single cell data has also raised new computational challenges. Unlike bulk expression datasets, that often profiled only a few samples, single cell datasets can contain hundreds, and even thousands, of expression profiles each [8, 35, 45]. In addition, single cell data is often noisier with potentiald ‘drop outs’ [29] making it harder to analyze. Consider for example one of the most exciting applications of single cell sequencing: the ability to identify and characterize new cell types and cell states [7, 30]. Recent work has used single cell expression profiles to discover new cells in developing lungs [42], new brain cells [7] and to refine several aspects of cell state transitions in differentiation studies [24, 17]. A key question that all such studies had to address is how to determine the similarity of the expression profiles of a pair (or larger sets) of cells? Another application for which the ability to compare single cell expression data between cell is critical is retrieval of similar cell types. Consider an experiment in which a population of cells taken from a diseased individual, or from a tumor, is profiled. One question that may be important for such analysis is to identify the specific types of cells that are present in the sample that was profiled, for example to determine which immune cells may have penetrated the diseased tissue [15]. While such analysis is often performed using markers, a much more comprehensive solution is to compare the various cell expression profiles to a set of curated single cells with known types.

In the above examples, comparisons or similarity analysis can either be performed using the measured expression values or after performing dimensionality reduction which may help reduce the noise associated with specific values. Indeed, several methods have been used and developed for performing such comparisons. The simplest, though one of the most popular, is based on principal component analysis (PCA). PCA has been used extensively for clustering single cells [5, 36, 45]. Other groups have developed new methods which build on PCA and extend it to improve clustering results. These include pcaReduce [48], which uses a novel agglomerative clustering method on top of PCA to cluster the cells. SNN-Cliq [47] constructs a *shared k*-nearest neighbor (KNN) graph over all the cells with the weight of each edge being the difference between *k* and the highest averaged ranking of the common KNN between two cells. It then tries to find maximal cliques in that graph in order to cluster the cells. ZIFA [29] develops a dimensionality reduction technique that takes into account the dropout characteristics of single cell sequencing data. SINCERA provides a pipeline for analysis of single cell gene expression data, one of whose tasks is to identify new cell types [14]. Clustering is done via hierarchical clustering with centered pearson correlation as the similarity measure. SIMLR [46] is another open-source tool that performs dimensionality reduction and clustering based on a cell similarity metric.

While PCA based approaches have been successful, they have so far been performed separately for each dataset. In contrast, for problems including retrieval we would like to obtain a reduced dimension for *all* cell types and experiments. In addition, PCA is an unsupervised method and so cannot find commonalities that are unique to specific types of cells but are less common in other types of cells. This latter sets can be very useful for grouping or retrieving single cell data.

Here we propose to replace PCA based dimensionality reduction with a supervised method based on deep neural networks. Neural networks are universal function approximators [16] and are very scalable in terms of training when using GPUs. These models use multiple layers, often with fewer nodes than the number of input values (genes), to integrate measured expression data. The networks are trained by maximizing their ability to identify the correct cell type from the values computed at the intermediate layers (Figure 1). Thus, the values computed for these intermediate layers can be used as a way to efficiently represent the input expression values. We tested various architectures for such networks, including architectures informed by prior biological data (such as protein-protein and protein-DNA interactions) and compared their performance to prior methods for analyzing single cell data. As we show, the learned networks captured several important biological aspects of the data. We observed improvements in clustering performance when using the neural networks computed values when compared to PCA, methods that use the measured expression data and prior methods for clustering single cell data. Finally, we show that the values obtained from the neural networks can improve the ability to retrieve the most relevant cells, for some cell types scientifically so.

**Figure 1:**
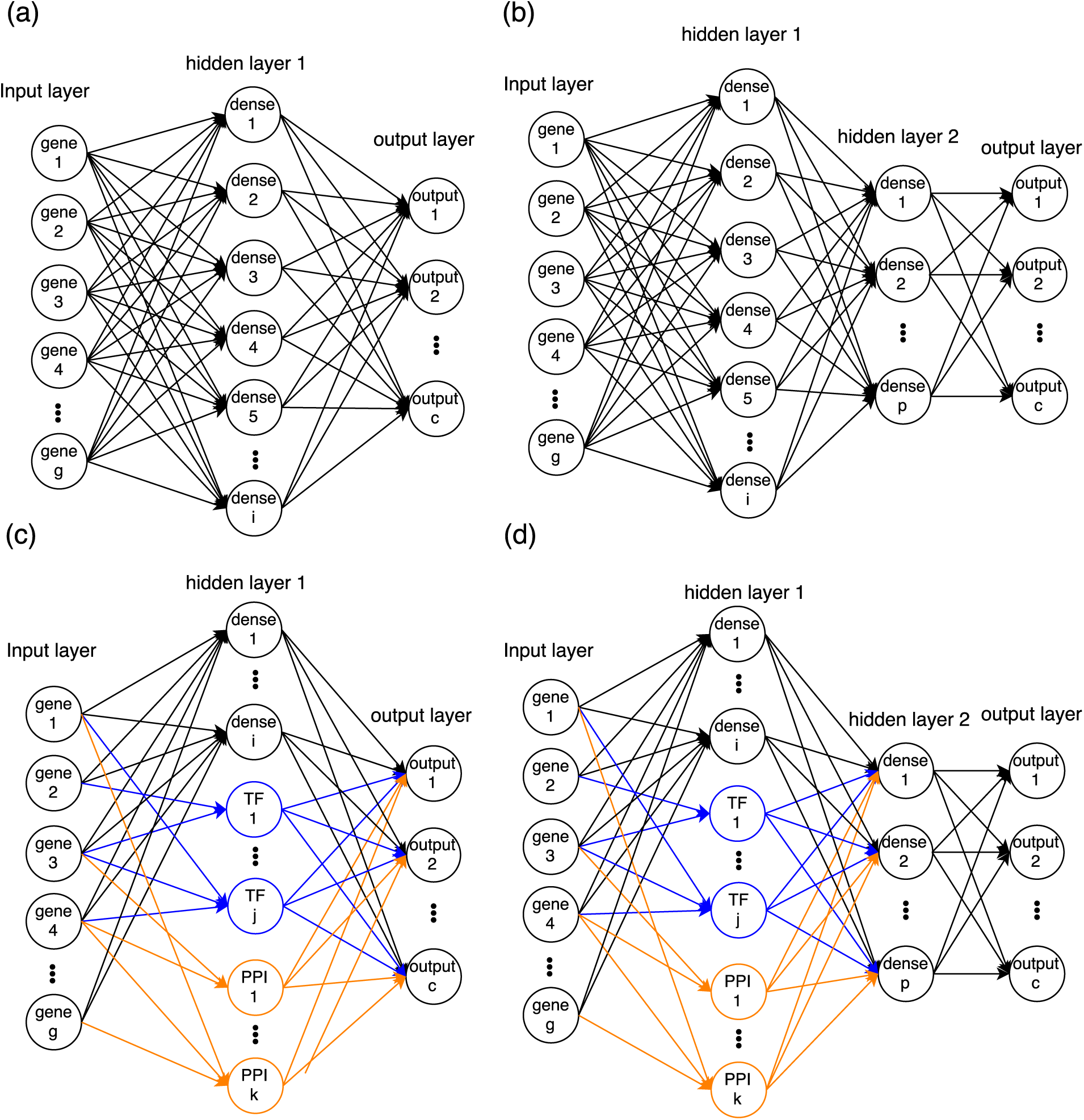
Network architectures of deep learning models used in this paper (a) Single-layered fully connected network (b) Two-layered fully connected network (c) Single-layered network with TF and PPI cluster nodes (connected only to their member genes) and fully connected dense nodes. (d) Two-layered network that are similar to the model in (c) but with an additional fully connected layer. Note that bias nodes are also included in each model.

## 2. Methods

### 2.1. Datasets used in our analysis

We collected a total of 33 datasets with more than 17,000 single cell expression profiles from published papers and from the Gene Expression Omnibus [3] (GEO). Supporting table and the supporting website provide full details on these datasets. We used 3 of these datasets which, combined, profiled 16 different types of cells and a total of 402 single cells for initial training, testing and evaluation of the method. We used 31 of the 33 datasets for the retrieval analysis (to avoid using different datasets from the same lab). All datasets are used in the retrieval application which is available from the supporting website. We curated all 33 datasets and assigned cell type labels to all single cell expression profiles.

### 2.2. Normalization and imputation

We tested a number of methods for normalizing data obtained from different labs and different platforms. We initially tested a novel normalization method for single-cell RNA sequencing data which is based on pooling across cells [26]. However, while results for the clustering analysis using this method were similar to the results presented below (see Supporting Table 3), the method required us to manually set several parameters (such as pool size) which made it hard to use for the larger retreival analysis. Instead, following prior work we normalized the data by converting all datasets to the the Transcripts Per Million (TPM) format [22, 37, 36, 35].

To combine protein-protein (PPI), protein-DNA (PDI) and single cell data (see below) we only used a subset of 9437 genes that were present in the 3 single cell training datasets and in the PPI and PDI datasets for training. For the much larger set of profiles used in the retrieval analysis, we used the same set of genes. Since these datasets were generated by different platforms and groups, counts for 2% of the genes were missing (on average) from each dataset. In order to use the NN method for analyzing these datasets we performed imputation for these missing genes as follows. Missing values for the retrieval analysis were first assigned the median gene expression value for the cell and then imputed with the average expression value for the k-nearest neighbor genes (we used k=10), where nearest neighbors were computed based on overall correlation [43]. Following TPM normalization and imputation, each gene was normalized to the standard normal distribution across samples since this is an essential step for NN training. We choose not to do the log-transformation because we found that it did not help the performance.

To account for the drop-outs in the imputation procedure, we tested a probabilistic model that would randomly assign values for 0 to a specific fraction *z* of the imputed genes instead of relying on the nearest neighbors for as discussed above. We tested several different values for *z* including 0 (no drop outs), 0.01, 0.03 and 0.05 using a cross validation strategy. Our results indicate that the best performance is achieved using *z* = 0 (see Supporting Table 4) and so this is what we used for the rest of the paper.

### 2.3. Protein-protein and protein-DNA interaction data

We used protein-protein (PPI) and protein-DNA interaction (PDI) data to determine the architecture of some of the NN we tested. We constructed a weighted, partially directed, protein interaction network using several databases including BIOGRID [39], HPRD [31] and also used Post-translational Modification Annotations from the HPRD. Protein-DNA interactions (PDI) were based on data from [34]. The data contained 160,000 PPI edges and 60,000 PDI edges between 16,671 genes and 348 TFs. The PPI data was weighted based on the different types of experimental evidence supporting each interaction [12].

### 2.4. A Neural Network representation of single cell expression data

We evaluated 4 types of neural network architectures [13] (Figure 1), and trained 5 models in total (detailed number of nodes for each model are shown in Table 1). All architectures include an input layer, one or two hidden layers (more hidden layers do not help the performance in the following experiments), and an output layer. The input layer encodes the expression values for the genes that are used in the analysis. The output layer encodes the probability of each cell type. Hidden layers are functions of the input and are used to efficiently encode the inputs such that the output can be the correct cell type given a set of expression values.

**Table 1:** The 5 different types of NN used in the paper

| No. | Model | layer1#node | layer2#node |
| --- | --- | --- | --- |
| 1 | Dense | 796 | X |
| 2 | Dense | 100 | X |
| 3 | Dense | 796 | 100 |
| 4 | PPI/TF+dense | 696+100 dense | X |
| 5 | PPI/TF+dense | 696+100 dense | 100 |

Note that No.1 and No.2 represent the same architecture (Figure 1 (a)) using different number of hidden layer nodes. The 696 in the PPI/TF models is from 348 TFs + 348 PPI groups. The number of nodes in the Dense models corresponds to the number of nodes in PPI/TF models for comparison. The additional 100 dense in each model are manually selected.

Specifically, we formulate our neural network model as follows. Let ***x***^(*i*)^ denote the output of i-th hidden layer. We use ***x***^(0)^ to represent the input of NN. To compute ***x***^(*i*)^, we perform forward propagation: (eq: 1)

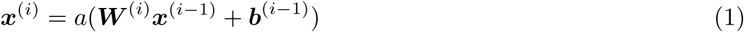

where *a* is the activation function, ***b*** is an intercept term and ***W*** is the weight matrix of each edges in the neural network. ***W*** and ***b*** are the parameters we need to learn.

We tested a number of possible activation functions including sigmoid, linear, relu, and tanh. Our analysis indicates that the hyperbolic tangent activation function (tanh) (eq: 2) leads to the best performance among these, and so we used it in the remainder of this paper. The tanh function is defined as:

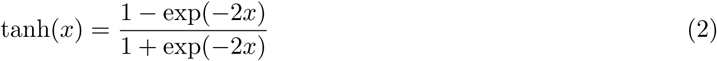

For the output layer, which performs discrete classification, we use the softmax activation function (eq: 3). Let ***x*** denote the input of the output layer (which is also the output of the last hidden layer), then we have the following:

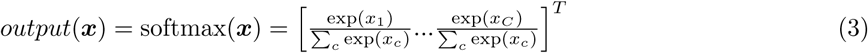

Where C encodes indices for all the cell types in training set.

The output of each node *c* in the output layer represents the probability *f* (***x***^(0)^)_*c*_ = *p*(*y* = *c|****x***^(0)^) that the input sample ***x***^(0)^ is obtained from that cell type. The loss function is categorical cross-entropy function (eq: 4):

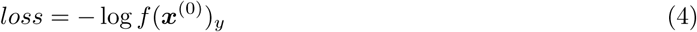

where *y* is the true label of the input ***x***^(0)^.

### 2.5. Architectures used in our NN method

While all networks used the same input and output layers, they differed in the structure and number of hidden layers. We tested models with 1 or 2 fully connected hidden layers (for the NN with 2 hidden layers, 796 nodes in the first hidden layer and 100 nodes in second hidden layer, Figure 1 (a)(b)). While these models have at most 2 hidden layers (more layers doesn’t help the performance of following experiments), the number of parameters that are fitted is very large (the architecture with the most parameters had 7.6 million parameters). Thus, even though these networks are not particularly deep, they require special hardware and software for training.

The above architectures do not make use of prior biological knowledge. To incorporate such information we also tested architectures that were based on PPI and PDI. Unlike the fully connected layers in the architectures described above, when using the biological data we connect the nodes in the first hidden layer to only a small subset of the input nodes. This greatly reduces the number of parameters that needs to be learned which can help with issues related to overfitting. Specifically, we have 348 hidden layer nodes based on the PDI data (one for each TF, connected only to genes regulated by that TF) and 348 nodes for the PPI. To divide the PPI graph into 348 subnetworks (where each subnetwork is connected to a node in the hidden layer) we used the ClusterOne [28] algorithm, which produces overlapping clusters so that, similar to the dense nodes, a gene can be connected to more than one PPI node. We also added 100 fully connected nodes (manually selected) to the hidden layer which account for missing data and TFs in the PPI and PDI. Still, the total number of parameters for this architecture is about 1 million, an order of magnitude less than for the fully dense architectures.

### 2.6. Unsupervised pre-training

The discussion above focused on supervised learning of NN (where the label is the cell type). In addition to supervised learning NN can also use unlabeled data, a practice that has proven useful in other domains [10, 4]. One kind of this NN is termed ‘autoencoder’ since the goal is to reconstruct the input layer values using a small number of hidden layer nodes (since the target is the input expression values no labels are needed to train such autoencoders). While unsupervised, autoencoders have been shown to successfully identify input combinations that affect the overall set of values. Given the large number of parameters in a NN the ability to train autoencoders and use the parameters learned as priors for a supervised learning procedure improves the initialization of the model and often leads to better generalization performance [10]. We have thus tested the use of 1-layer denoising autoencoders (DAE) when testing the method on large datasets (retrieval datasets). We train DAE to reconstruct the original input from corrupted input with the noise sampled from a standard normal distribution multiplied by a noise level of 0.1. The architecture of DAE is similar to Figure 1 (a) except that the output layer is changed to be the reconstructed input. All layers in the DAE use the tanh activation function and mean square error as the loss function. Here we used 100 and 796 nodes in the hidden layer of the DAE, similar to the numbers used for the supervised models to make sure that the weight of DAE can be used as pre-trained weights of supervised models.

### 2.7. Learning model parameters

All models were implemented in the Keras tool [6] with some modifications to accommodate the sparse layer connections for TF and PPI nodes. The models are trained using a stochastic gradient descent optimizer with a learning rate of 0.1, decay 10^-6^, momentum 0.9, and Nesterov accelerated gradient. These parameters are manually selected for convergence. We used 100 iterations (which were manually selected and enough for reaching convergence) to train each model with a mini-batch size of 10, which is the number of samples to fit at each iteration. Detailed information about the dimensions of the different architectures is provided in Table 1. It took us 30 seconds to train the largest supervised NN (7.6 million parameters, 402 cells), 40 minutes to train the largest unsupervised NN (15 million parameters, 17000 cells) on a machine with 4 Intel(R) Xeon(R) CPU E5-2620 v3 (2.40GHz each core, 24 cores in total), 4 Nvidia GTX 1080 GPUs and 128 GB RAM.

### 2.8. Biological analysis of learned models

To determine the biological relevance of the parameters learned from the NN we analyzed significant gene groups for each cell type in the PPI/PDI model (Figure 1 (c)). For this we identify the top 10 most highly weighted (hidden layer) nodes for each output layer node (corresponding to the different cell types). Some of the selected nodes are explained by the TF or the PPI they represent. For the other (100 nodes in the hidden layer initially connected all input genes) we perform GO analysis based on the set of input genes that are connected to these nodes with a high (absolute value) weight. We used gprofiler [32] to perform GO analysis because it provides command-line interface for access.

### 2.9. Comparisons to prior clustering and dimensionality reduction methods

To perform dimensionality reduction based on the NN results we extract the values computed by the last hidden layer of each architecture we tested. We next use a simple clustering method (K-means++ clustering [1]) to perform unsupervised grouping of cells using a test set (not used in the NN training) and the results are compared to prior methods suggested for clustering single cell expression data. For such comparisons we perform experiments in which we leave out 2, 4, 6, or 8 random cell types of the 16 types in our analysis set. We next cluster the left out data using the reduced representation obtained from the last hidden layer of the NN and use the adjusted random index (ARI) to compare the clustering results with the true labels. ARI counts the number of agreements and disagreement between two groupings while adjusting for random performance into account. It is defined as follows. Let *X* = {*X*_1_*, X*_2_*, …, X*_*r*_}, *Y* = {*Y*_1_*, Y*_2_*, …, Y*_*s*_} be two groupings. We can summarized the overlap between *X* and *Y* using a table *N* where *N*_*ij*_ = *|X*_*i*_ ⋂ *Y*_*j*_ |is the number of objects in common. Let *a*_*i*_ =∑_*j*_ *N*_*ij*_, *b*_*j*_ = ∑_*i*_ *N*_*ij*_, *n* be the total number of samples, then we set:

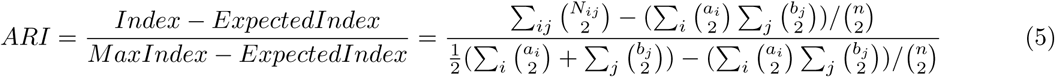

We repeat the experiment for 20 times with fixed random seeds 0-19, each time with different random left out cell type and obtain an average ARI value.

### 2.10. Cell retrieval method

To measure the performance of cell retrieval, the percentage of the desired cell type in the top k nearest neighbors (here we used k=100 is calculated for different single cell expression data representations (NN, PCA, measured values). To reduce the ability of the retrieval analysis to rely on artifacts for correctly identifying the cell types (for example, experiments from the same lab) we only perform this analysis on cell types that were profiled in *different* datasets out of the 31 used for this analysis. We thus used 9 different cell types for this analysis though the database itself contains over 100 cell types (including subtypes). Cell types that are not selected can thus still be in the retrieved set of k nearest cells which makes the analysis more similar to how we envision the usage of such database. We use the mean of average precision (AP) to evaluate the retrieval performance. AP corresponds to the area under the precision-recall curve for the retrieved cells. The final result for all cell types is a weighted mean so that every dataset (cell type) has equal weight.

## 3. Results

We learned parameters for the different NN architectures discussed in Methods and used the resulting models to test the method and to compare it to prior methods for dimensionality reduction of scRNA-Seq data and for clustering and comparing such data.

### 3.1. Testing and comparing the NN method

To test our method and to compare it to other methods for clustering single cell time series expression data we used 3 single cell expression datasets with 16 types of known labels (See supporting table 6 for complete list). Data was downloaded mostly from GEO and processed as discussed in Methods (section 2.1). We identified 9437 genes that had expression values in all training datasets and used them to learn the (deep) NN using various architectures (section 2.4). For each architecture, input values where directly connected to one or more hidden layers and the output layer encoded the label for each of the datasets. Thus, the goal of the NN was to identify a reduced dimension representation of the expression values (where the number of nodes / values is a function of the specific architecture) that leads to the most accurate assignment of cell type for each dataset. Training and testing accuracy of all NN models reached nearly 100% (while this was a challenging multi-label classification task with up to 16 possible labels the number of parameters is very large as mentioned in Methods which can explain the good performance on the training set). See supporting table 6 for the full list of the 16 cell types.

Given the ability of the method to accurately assign training data we next asked how well the resulting representation can be used on test (unobserved) data. For this we performed a number of different analyses in which we divided the single cell datasets using subset to learn the model and the rest to test it. For testing, we compared the ability of a simple clustering method (k-means++ algorithm with k representing the known number of left out cell types) to accurately sub-divide test data according to their cell types. We first construct the deep NN using the portion of cells used for training. Next, for each of the test cells we run them through the NN and extract the values of the smallest hidden layer (depending on the architecture) and use these vectors in the clustering procedure. Thus, clustering is based on gene expression combinations determined by the NN. We also note that the training and test sets are comprised of different cell types and so this is not a supervised classification task. Instead, the goal is to see if the parameters learned using the labeled data can improve the *unsupervised* analysis of the other cell types. We also calculate the performance of pretrained models, but the result are similar to the model without pretraining. Therefore, we put the results of clustering performance with pretraining into supplementary table 5.

We used the clustering results to compare the NN based analysis to a number o unsupervised clustering and dimensionality reduction methods that have been proposed for single cell analysis. Specifically, we compared our method to PCA, pcaReduce, SINCERA, SIMLR and SNN-Cliq. For the PCA based methods we tested both 100 and 796 components, similar to the dimensions of the output of our method. PcaReduce generates hierarchical structure of clustering based on PCA reduced dimensions. SINCERA provides a pipeline of analysing single cell data, including clustering. For clustering, SINCERA uses hierarchical clustering, tight clustering [44] or consensus clustering [27]. We only show the result of hierarchical clustering here because it is the default setting of SINCERA and the other clustering methods often generated error messages when applied to our dataset. SNN-Cliq uses shared nearest neighbors (SNN) to define the similarities between cells and then clusters them using graph based methods. SNN-Cliq sometimes generates error messages when the number of cell types (*k*) is 2, so we left it out in this comparison. We also tried to compare our results to ZIFA. However, ZIFA did not finish after running for two days when trying to cluster 300 cells with 9437 expression values each. To improve its run time we reduced the number of genes to 1356 (selecting only genes that have non-zero values in 90% of samples) but the performance of ZIFA on this data was not good (much lower than the results presented in the comparison table) and so we left it out. As can be seen in Table 2, the clustering results of all the different architectures we tested outperformed all other methods including PCA, SIMLR, SNN-Cliq and using the original (non reduced) set of expression values (see also Supporting Figure 1). One way to explain this result is the fact that unlike other methods our method learns the best way to represent a reduced dimension for single cell data (even though the comparison is based on an unsupervised clustering task and learning is done on a completely different set of experiments and cell types) whereas the other methods are fully unsupervised. While all NN architectures performed better than other methods, the simplest NN architecture we tested (1 fully connected layer with 796 hidden nodes) performed best, though not by much. However, as we show below, when testing on a much larger set of data the biologically motivated architectures do better than the fully dense ones.

**Table 2:** Average adjusted random index (ARI) for 20 clustering experiments (using different random initializations

| feature \ # testing cell type | 2 | 4 | 6 | 8 | Average |
| --- | --- | --- | --- | --- | --- |
| original | 0.943 | 0.687 | 0.651 | 0.613 | 0.723 |
| PCA 100 | 0.943 | 0.582 | 0.596 | 0.554 | 0.668 |
| PCA 796 | 0.943 | 0.691 | 0.600 | 0.549 | 0.696 |
| pcaReduce | 0.760 | 0.692 | 0.677 | 0.683 | 0.703 |
| SIMLR | 0.850 | 0.749 | 0.663 | 0.625 | 0.722 |
| SNN-Cliq | X | 0.716 | 0.661 | 0.435 | 0.604 |
| SINCERA hierarchical clustering | 0.980 | 0.797 | 0.738 | 0.674 | 0.797 |
| SINCERA tight clustering | X | X | X | X | X |
| SINCERA consensus clustering | X | X | X | X | X |
| Dense 1layer 100 | 0.978 | 0.882 | 0.755 | 0.656 | 0.817 |
| Dense 1layer 796 | 0.981 | **0.885** | 0.765 | **0.708** | **0.835** |
| Dense 2layer 796/100 | **0.984** | 0.880 | 0.763 | 0.630 | 0.814 |
| PPI/TF 1layer 696+100 | 0.981 | 0.839 | **0.781** | 0.665 | 0.817 |
| PPI/TF 2layer 696+100/100 | **0.984** | 0.876 | 0.752 | 0.662 | 0.818 |

### 3.2. Functional analysis of hidden layer nodes

While NN are often treated as “black boxes” following recent studies we attempted to characterize the input and hidden nodes (genes / TFs) that had the highest impact on the ability to accurately classify various cell types [40, 38]. Such analysis can provide both, functional interpretation of the parameters that are learned for the method as well as characterization of the different types of genes and their regulators that are most relevant to specific types of cells. To perform such analysis, we analyze the top 10 most highly weighted groups (hidden nodes) for each cell type with the NN in Figure 1(c). To analyze the groups we either used the known characteristics of the node (since, as mentioned in Methods (section 2.8) some of the nodes represent groups of genes known to be co-regulated by the same TF or to be interacting (PPI)) or GO analysis on fully connected nodes to determine the function of genes associated with that node. Table 3 presents results for a subset of the cell types and nodes which are manually identified. As can be seen, the top scoring nodes were often very relevant for the specific functions that are performed by the cells or the TFs that regulate these functions. For example, the learned NN correctly associated genes related to proliferation and differentiation with ES cells while nodes that scored high for immune response categories were mainly associated with Bone Marrow-derived Dendritic Cells (BDMCs). See also Supporting Website for tables summarizing more than 1000 significant GO categories for combinations of cell types and nodes based on this analysis.

**Table 3:** Examples of highly ranked nodes for some of the cell types used for learning of the NN.

| cell type | TF/PPI/Dense node | Corrected p-value | GO function / reference |
| --- | --- | --- | --- |
| stem cells (ES) | dense 35 | 3.25E-07 | cell differentiation |
| stem cells (ES) | dense 24 | 4.44E-15 | Factor: E2F-3; [20] |
| stem cells (ES) | dense 24 | 5.78E-12 | Factor: IRF6; [2] |
| stem cells (ES) | dense 24 | 2.53E-08 | system development |
| BMDC | dense 10 | 1.56E-05 | immune system process |
| BMDC | dense 67 | 1.59E-11 | positive regulation of immune system process |
| BMDC | dense 36 | 2.06E-06 | response to cytokine |
| fibroblast | ppi 223 | 3.16E-06 | Focal adhesion [33] |
| fibroblast | ppi 301 | 2.85E-20 | acetyltransferase complex [11] |
| zygote | ppi 10 | 6.85E-07 | regulation of cell proliferation |
| zygote | ppi 280 | 7.36E-09 | cell junction |
| zygote | TF: foxd3 | X | TF: foxd3 [9] |

Some of the nodes were based on TF-gene interactions and thus represent a specific TFs (Foxd3). For these we rely on the function of the TF to characterize the node in the table. Other nodes are either based on PPI (for example PPI223) or on groupings learned by the algorithm (for example, dense24). For these we performed GO analysis on the set of highly ranked genes for these nodes. Several other relevant TFs and functions were found for other cell types as well. See Supporting Website for complete list.

In addition to the analysis of connection weights for the NN that were learned using labeled data, we also analyzed the values for the nodes obtained by the pre-trained models. Recall that for the pre-training we are using a fully unsupervised approach where the goal is to reconstruct the input data using a small set of highly connected nodes (100 in our case). We hypothesized that some of the values learned for these nodes may actually reflect key genes that either regulate other genes (and so their values are enough to reconstruct the full expression set) or important groups of co-expressed genes. We thus selected, for each of the 100 dense nodes in the pre-trained mode, the set of 3 most similar genes based on Pearson correlation. The results are presented in Supporting Table 7. As can be seen, many of these genes are ribosmal genes which are indeed a large and coherently expressed group that is captured by some of the nodes. GO analysis of the selected nodes (Supporting Table 8 shows part of the results.) indicated that a a significant category for these is ‘nucleic acid binding’ (corrected p-value = 9 *\** 10^-9^) indicating that the model captures some of the TFs that are likely regulating the expression in the different single cell types. See Supporting Results and website for complete details.

### 3.3. Retrieval of single cells based on expression

In most single cell studies, hundreds or even thousands of cells are being profiled. In almost all cases, including cancer [22], brain studies [18, 49, 25, 21] and more, several different types of cells are profiled in the same experiment. In some cases these cells can be characterized using marker genes and assigned to a specific cell type. However, in most cases at least some of the cells cannot be fully assigned since either they do not contain any of the known markers or they contain several subsets of such markers. In most cases researchers classify such cells using clustering allowing them to identify several groups within the sampled cells. However, such analysis is unsupervised and so its not always clear what each of the clusters corresponds to. Identifying the composition of cells is important, for example in cancer studies where notable differences between outcomes have been attributed to the amount of immune cells that are present in the tumor.

Thus, an important problem in single cell expression analysis is assignment. One way to address this problem is to compare uncharacterized cells to cells that have already been characterized by prior experiments either using follow up studies or because of their known origin. To enable such analysis we collected 31 single cell expression datasets with more than 17000 samples and created a database in which we stored both the expression measurements for these cells as well as the assignment for the cell from the paper / experiment in was profiled in (if available). Using such database we can find, for each new cell profiled in an experiment, the most similar cells in the database and based on their annotations, annotate the uncharacterized cells.

A key issue for implementing such strategy is how to identify the most similar cells in the database when given a new query cell. A simple solution is to use the expression values themselves. However, such method is both inefficient (requiring us to store the actual values for all genes in the database as well as perform a large number of operations for each pairwise comparison to compare all 20K+ genes). An alternative, which is more efficient and may be more robust to noise is to first reduce the dimension of the database data and the query cell and then perform the retrieval search in the reduced dimensional space. Such reduction can be done either using PCA or using the NN approach discussed above. By using a lower dimension representation we drastically cut the storage requirements (by 91% to 99% depending on the architecture used) and the query run time. Specifically, for the 14000 queries we performed on the 17000 cells in the database we reduced the run time from 25 minutes (when using the observed expression values for 9437 genes) to less than 5 minutes when using the 796 features obtained from the NN. The datasets of the query cell types are listed in Supporting table 2.

In addition, and most importantly, such reduced dimension greatly improves performance. To test various ways of querying single cell expression data we held out complete datasets for which we had a similar dataset in the database from a different lab / paper. This helps ensure that results are not effected by unique lab methods but are rather related to biological similarities. We identified 31 different datasets that, combined, profiled close to 14K cells. For each of these held out datasets we searched for the most similar cells in our database. As before, we compared the NN results to results obtained when using the measured expression values and PCA with 100 or 796 dimensions. As for the NNs, since we are dealing with much more data compared to the clustering analysis above we tested two NN variants. The first is similar to the one described above for clustering while the second is based on using DAE to initialize the parameters of the network (using unlabeled data) followed by supervised learning as discussed in Methods.

To evaluate the different methods, for each query cell we identify the top *k* most similar cells in the database (where similarity is either based on Euclidean distance for all genes or for the reduced dimension representation obtained by each method). We use *k* = 100 in this paper, though results are similar when using *k* = 10. We next use the top *k* matches to compute the mean average precision (MAP) for the correct cell type for each query (Methods).

As can be seen in Table 4, all methods that relied on reduced dimensions did much better than the method that used the measured expression values (20% average improvement for PCA and almost 40% improvement for some of the NN methods when compared to using the measured expression values for all genes). Comparing the reduced dimensionality methods themselves, we observe that NN with more hidden layers (in our case 2) are doing, on average, better than NN with a single hidden layer indicating that non linear relationships may be important for characterizing single cell expression. These multi-hidden layer NN are also performing better than PCA.

**Table 4:** Average retrieval performance across the different cell types

| models | HSC | 4cell | ICM | spleen | 8cell | neuron | zygote | 2cell | ESC | mean |
| --- | --- | --- | --- | --- | --- | --- | --- | --- | --- | --- |
| Original | 0.081 | 0.361 | 0.058 | 0.987 | 0.279 | 0.372 | 0.468 | 0.556 | 0.705 | 0.430 |
| PCA 100 | 0.299 | 0.508 | 0.01 | **0.996** | 0.351 | 0.646 | 0.539 | 0.616 | 0.722 | 0.521 |
| PCA 796 | 0.227 | 0.548 | 0.022 | 0.994 | 0.242 | 0.622 | 0.419 | 0.642 | **0.833** | 0.505 |
| DAE 100 | 0.236 | 0.411 | 0.016 | 0.973 | 0.503 | 0.628 | 0.193 | 0.728 | 0.544 | 0.470 |
| DAE 796 | 0.14 | 0.423 | 0.035 | 0.992 | 0.399 | 0.692 | 0.399 | **0.743** | 0.432 | 0.473 |
| Dense 1layer 100 | 0.102 | 0.662 | 0.038 | 0.953 | 0.739 | 0.485 | 0.522 | 0.717 | 0.604 | 0.536 |
| Dense 1layer 100 pretrain | 0.23 | 0.49 | 0.153 | 0.984 | 0.463 | 0.64 | 0.408 | 0.734 | 0.532 | 0.515 |
| Dense 1layer 796 | 0.082 | 0.599 | 0.096 | 0.988 | 0.472 | 0.389 | **0.563** | 0.732 | 0.683 | 0.512 |
| Dense 1layer 796 pretrain | 0.116 | 0.515 | 0.073 | 0.991 | 0.463 | **0.702** | 0.423 | 0.714 | 0.446 | 0.494 |
| Dense 2layer 796/ 100 | 0.069 | 0.77 | 0.065 | 0.956 | 0.896 | 0.563 | 0.275 | 0.673 | 0.583 | 0.539 |
| Dense 2layer 796/ 100 pretrain | 0.164 | 0.648 | 0.035 | 0.987 | 0.715 | 0.633 | 0.498 | 0.747 | 0.507 | 0.548 |
| PPITF 1layer 696+100 | 0.078 | 0.636 | 0.148 | 0.965 | 0.667 | 0.464 | 0.202 | 0.314 | 0.63 | 0.456 |
| PPITF 1layer 696+100 pretrain | 0.168 | 0.557 | 0.028 | 0.982 | 0.55 | 0.647 | 0.447 | 0.665 | 0.569 | 0.513 |
| PPITF 2layer 696+100/ 100 | 0.068 | **0.771** | 0.182 | 0.956 | **0.849** | 0.561 | 0.415 | 0.553 | 0.71 | 0.563 |
| PPITF 2layer 696+100/ 100 pretrain | **0.397** | 0.614 | **0.185** | 0.975 | 0.725 | 0.626 | 0.435 | 0.688 | 0.554 | **0.578** |

We also observe that both, the use of prior biological knowledge to define the NN architectures (PPITF networks) and the use of pre-training using DAE improves the overall accuracy of the retrieval. Specifically, the best performing method (achieving an improvement of more than 11% over PCA) is the PPITF 2layer 696+100 / pretrain which combines all these features (2 layers, pre-training and the use of prior knowledge). Other architectures that use prior knowledge are also better than their dense counterparts. In contrast, DAE on their own (4th and 5th rows) are not as effective as supervised models and so they are probably best for initializing rather than for final model selection.

## 4. Discussion and future work

While single cell analysis holds great promise, it also raises new questions. Given the number of cells that are profiled in each experiment, which can reach thousands [49, 23], new methods are required for accurately and efficiently clustering this data while overcoming issues related to the stochastic nature of gene expression even in similar cells, noise and missing values. A related problem is the ability to compare expression profiles from cells so that cell type assignments can be determined not just based on a few marker genes but rather based on the overall expression of all genes in the cells profiled.

In this paper we developed and tested solutions based on deep neural networks for these problems. The advantage of such networks is that they can learn the importance of different combinations of gene expression levels for defining cell types and such combination are usually more robust than values for individuals genes or markers. We tested a number of different activation functions for this data and several NN architectures, including architectures that are constrained by prior biological knowledge. As we show, the NN achieve very good classification performance on training data and improve upon prior methods when used to cluster datasets from experiments that were not used in the training. We also performed functional analysis of the set of highly weighted nodes for each cell type and showed that even though NN are often described as a black box learning method, many of these are functionally related to the cell type they were selected for.

As a final application we used the reduced representation obtained form the NN to query a large database of over 17K single cell expression data in order to determine the cell type of a newly profiled single cell. As we show, using such representation greatly improved the performance of the retrieval analysis while reducing the overall runtime and storage required. The Supporting Website provides an implementation of the retrieval method which can be used by researchers to determine cell types for newly profiled single cells.

While the results are encouraging, there are several directions for future work which we would like to explore. These include testing more involved (deeper) architectures, integrating additional types of prior biological knowledge into the model and an automated tool that can download new single cell expression data in order to increase the set used by the retrieval application. A major challenge with the latter direction is the ability to automatically assign cell type from published expression data given the various ways in which people define and encode such information.

